# What group averages conceal: functional heterogeneity in human eyeblink habituation

**DOI:** 10.64898/2026.06.21.733594

**Authors:** Omar David Perez, Natalia Cancino, Daniel Hermosilla, Fabian A. Soto, Edgar H. Vogel

## Abstract

In animal learning research, learning is often represented by plotting a behavioral measure as a function of training trials. A particularly clear case is habituation, a basic form of learning in which repeated presentation of a stimulus produces a decrement in responding. Although retention tests provide the strongest basis for evaluating durable habituation once short-lived performance effects have dissipated, the pattern of response change across stimulus repetitions, or habituation curve, remains theoretically and empirically relevant because it is used to characterize determinants of habituation, individual and clinical profiles, and functional forms, including linear, curvilinear, asymptotic, and mixed incremental–decremental patterns of responding. However, group averaged curves may conceal substantial individual heterogeneity. Here, we analyzed archived human eyeblink habituation data from 157 participants to ask whether the curve shape selected for the group average reflects the curve shapes observed at the individual level. Five candidate functions were fitted separately to each participant and to the corresponding group average. No single function characterized most individuals. More importantly, the model selected for the group average differed from the most frequent individual model in all four groups. When data were pooled across groups, the average favored a dual-process form, a shape that matched the individual plurality in none of them. Simulation analyses showed that averaging heterogeneous individual trajectories can itself produce a group curve that favors a more complex model. Our findings show that group averaged habituation curves should not be treated as direct descriptions of the typical individual trajectory.

## Introduction

A long-standing practice in behavioral research is to characterize psychological processes from the shape of group-average curves. When responding changes systematically across trials, researchers average across participants, fit parametric functions to the resulting curve, and interpret the winning model as a description of the underlying individual process. This convention is partly computational—because fitting nonlinear functions to noisy individual sequences is demanding—and partly inferential: the group average is assumed to represent the most common individual trajectory.

What is less clear is whether that inference is warranted when individuals differ in their process parameters. The problem was identified early by Estes (1956) in the context of learning curves. Estes showed that when individuals differ in a rate parameter, the group average does not generally have the same form as any individual curve. For example, a group of subjects who each follow simple exponential declines with different rates can yield an average that is shallower, slower, and better described by a more complex model. This is not merely a sampling issue; it follows from averaging nonlinear functions across a distribution of parameters. Ashby et al. (1994) formalized the conditions under which such distortions occur, and Lewandowsky (1995) showed that the same logic applies to forgetting functions. The same concern arises whenever individual trajectories are summarized for modeling or visualization (Farrell and Lewandowsky, 2018). Together, these analyses suggest a simple methodological principle: conclusions about individual curve shape should not be drawn from the group mean alone.

Gallistel et al. (2004) provided a direct demonstration of this point in the context of learning. They argued that individual acquisition curves in conditioning are better described by abrupt, step-like transitions—responding near zero until a critical trial, then jumping to asymptote—than by the gradual error-correction curves that are typically reported. When subjects differ only in *when* the step occurs, the group average smooths across those transitions and produces a gradual-looking curve that no individual exhibits (Figure 1). The apparent continuity is an artifact of averaging heterogeneous step locations, not evidence of a gradual underlying process.

**Figure 1:**
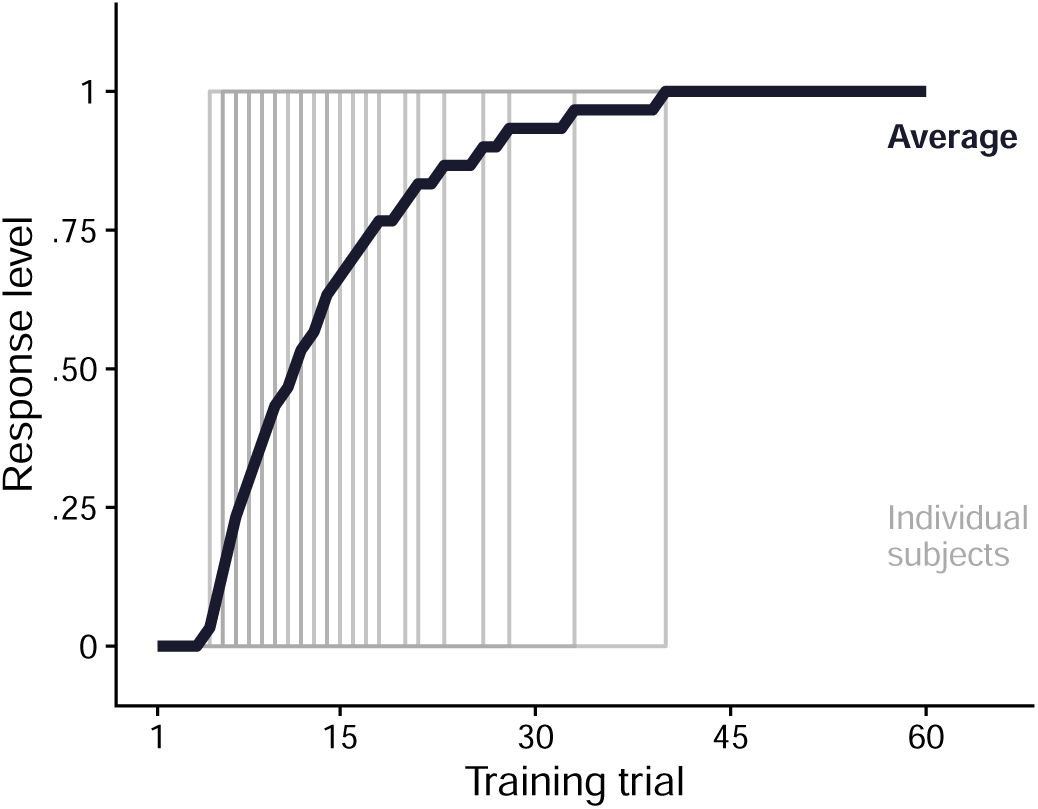
Schematic of the Gallistel et al. (2004) argument. Each grey line is one simulated subject whose response jumps from zero to asymptote at a different training trial. The black curve is the group average. No individual shows gradual learning; the smooth average is an artifact of averaging heterogeneous step locations.

One domain where this practice is particularly common is habituation research. Habituation is a decrease in responding produced by repeated stimulation, and is among one of the most conserved behavioral phenomena across species (Thompson and Spencer, 1966; Rankin et al., 2009; Thompson, 2009). The trajectory of response change across repetitions has been treated as a central feature of habituation: Rankin et al. (2009) noted that response decrement is often exponential but may also be linear, and that some responses may show facilitation before decrementing. This trajectory can be characterized by several components, including the initial response level, rate and form of decrement, possible increments, and asymptotic level. These components have direct relevance to clinical comparisons: studies of anxiety, schizophrenia, and related conditions routinely characterize patient groups by the shape of their habituation trajectory (McDiarmid et al., 2017; Cancino et al., 2026).

The most influential account of such response trajectories is the dual-process theory of Groves and Thompson (1970), which assumes that repeated stimulation engages two opposing processes in parallel—habituation, which decreases responding, and sensitization, which increases it. The net output can produce an initial rise before decline, the characteristic inverted-U pattern reported in some species and stimulus conditions (e.g., Peeke and Veno, 1973; Petrinovich and Peeke, 1973). Related analyses of defensive responding further show that repeated aversive stimulation may generate both response-decremental and response-potentiating tendencies (Wagner and Vogel, 2010), which means inter-individual variability in curve shape is expected, not exceptional. Because dual-process theory makes specific predictions about curve shape, the standard practice has been to fit these models to the group-average curve.

Whether group averages distort the functional form of individual habituation curves has not been directly tested. This question is related to a broader concern in learning theory about the interpretation of acquisition and habituation curves. Several authors have warned that response curves can confound learning with performance, because changes observed during training reflect not only what has been learned, but also the conditions under which that learning is expressed and measured (Hearst, 1988; Rescorla, 1988; Rescorla and Holland, 1976). In the specific case of habituation, this concern has motivated proposals to rely more heavily on common-test or retention-test procedures, rather than inferring the strength or amount of habituation directly from the response decrement observed during repeated stimulation (Colwill et al., 2023; Davis and Wagner, 1968). Nevertheless, abandoning curves altogether would also discard an important feature of the phenomenon. The trajectory of responding across repeated presentations captures the short-term expression of habituation, including transient response dynamics that may reflect short-term memory for recent stimulation and stimulus processing, even when they do not provide a pure measure of durable learning (Jorquera et al., 2025). The present study therefore focuses on a narrower but important question: when habituation curves are used to characterize performance across trials, does the functional form selected for the group average correspond to the functional forms observed in individual subjects? When individuals differ in their decay dynamics, the group mean can select a more complex model than the one that fits most participants. McDiarmid et al. (2017) called for individual-level analysis as a precondition for comparing habituation across populations and disorders, and recent reviews from the same laboratory as the data reanalyzed here have highlighted exactly this problem (Becerra et al., 2026; Cancino et al., 2026). The present study asks a direct question: when the same candidate models are fit to individuals and to the group average within the same experiment, does the winning model for the average match the individual plurality?

To address this question, we reanalyzed archived human eyeblink habituation data from three previous studies conducted by our research group at the University of Talca (*N* = 157 across four Experiment × ITI cells). We fit five candidate models to each subject and to the corresponding group-average curve. We then compared the winners within each group, repeated the comparison after pooling datasets under different weighting rules, and asked whether heterogeneous decay rates or missing trials could account for the observed pattern.

## Method

### Data set and analytic overview

The data analyzed in the present study were drawn from three previous human eyeblink habituation experiments. Across these experiments, participants received repeated presentations of a stimulus capable of eliciting an eyeblink response. The experiments varied in the stimulus used, the response measures recorded, the number of trials, the intertrial interval (ITI), and the presence of additional sessions or experimental manipulations.

For the purposes of the present analysis, we selected participants whose relevant habituation phase involved no manipulation other than the repeated presentation of the eliciting stimulus. Within those participants, we analyzed only the first uninterrupted sequence of trials delivered at a constant ITI. Thus, the analyzed data were restricted to trial sequences that provided a continuous record of responding to repeated stimulation, before any additional experimental manipulation was introduced. This selection criterion allowed the same curve-fitting procedure to be applied to each participant and to each group average.

Because one of the three experiments included two ITI conditions, the final dataset comprised four Experiment × ITI cells, with 157 participants in total (Table 1). The primary analyses were based on single-trial eyeblink amplitudes measured by an infrared LED sensor that detected eyelid closure. Trials with missing or nonpositive amplitude values were not entered into the analyses. Amplitudes were analyzed on their observed raw scale, without *z*-scoring or between-subject normalization. EMG data available for two of the experiments were analyzed separately as a replication and are reported in Appendix B. Additional protocol details for each experiment are provided in Appendix A.

**Table 1:** The four LED Experiment × ITI cells included in the present analysis.

| Cell | Study | ITI (s) | Trials | $N$ |
| --- | --- | --- | --- | --- |
| 1 | 1 | 30 | 100 | 50 |
| 2 | 2 | 8 | 50 | 67 |
| 3 | 3 | 10 | 50 | 20 |
| 4 | 3 | 30 | 50 | 20 |

The question at issue in this study was whether the curve family selected for the group average also corresponded to the curve family most commonly selected at the individual level within each Experiment × ITI cell. We proceeded in three stages, ordered by inferential priority. First, we analyzed each Experiment × ITI cell separately. Second, we repeated the comparison after pooling data across experiments under predefined averaging and weighting rules. Third, we conducted robustness checks to evaluate whether the observed pattern could be attributed to heterogeneous decay rates, missing trials, or the model selection procedure. The same trial-handling rules, model-selection criterion, and candidate curve models were used throughout. All the models are described in the following sections.

### Apparatus and dependent variable

Across the three experiments, eyeblink magnitude was recorded with head-mounted infrared LED sensors on the Eyeblink Conditioning System (EBC) apparatus (Ponce et al., 2015). The signal was sampled at 1 kHz and low-pass filtered at 40 Hz offline. Each trial was reduced to a single amplitude: the recording was windowed from 500 ms before stimulus onset to 300 ms after onset, prestimulus baseline activity was subtracted, and blink magnitude was taken as the maximum value in the post-onset window after baseline correction. This single per-trial amplitude is the dependent variable throughout.

### Time axis

Within each Experiment × ITI cell, raw trial number *t* was mapped to standardized session progress,

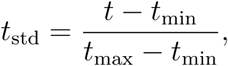

where *t*_min_ and *t*_max_ are the smallest and largest trial numbers present in that cell — that is, the first and last trial of the session — and are therefore identical for every participant in the cell. The first trial of the session maps to 0 and the last to 1, regardless of which trials a given participant happens to have retained.

This was a deliberate choice. Fixing the endpoints by the session rather than by each participant’s own retained range preserves the absolute position of every trial within the session. A participant who is missing late trials does not have their last retained trial stretched to 1; instead they occupy only the portion of [0, 1] corresponding to the trials actually observed. For example, in Study 2 (50 trials) a participant with valid data on trials 1–48 spans [0, 47*/*49], and a participant whose first valid trial is trial 5 begins at 4*/*49 rather than at 0. The transformation therefore retains information about where missing trials fall in the original sequence, which is what makes the missing-trial analysis reported below meaningful. Rescaling each participant to their own first and last retained trial would instead have discarded that absolute temporal position and stretched truncated sequences to fill [0, 1], making the position of missing trials unrecoverable.

The same standardized axis was used for every participant and for the group-average curve. This ensured that rate parameters in the candidate functions described below were expressed on a common relative-session scale, rather than on study-specific trial counts. For example, a rate parameter refers to change across the analyzed sequence from session start (0) to session end (1), regardless of whether the original cell contained 50 or 100 trials. The transformation does not mix data across studies; each cell is rescaled using only its own trial range. All models were fit as functions of *t*_std_; for brevity, *t* denotes *t*_std_ throughout.

### Candidate curve models

Five parametric families were fit to each trial series, with *t* as the independent variable and LED amplitude as *y* (Table 2; Figure 2). Nonlinear rate and amplitude parameters were constrained to be nonnegative.

**Figure 2:**
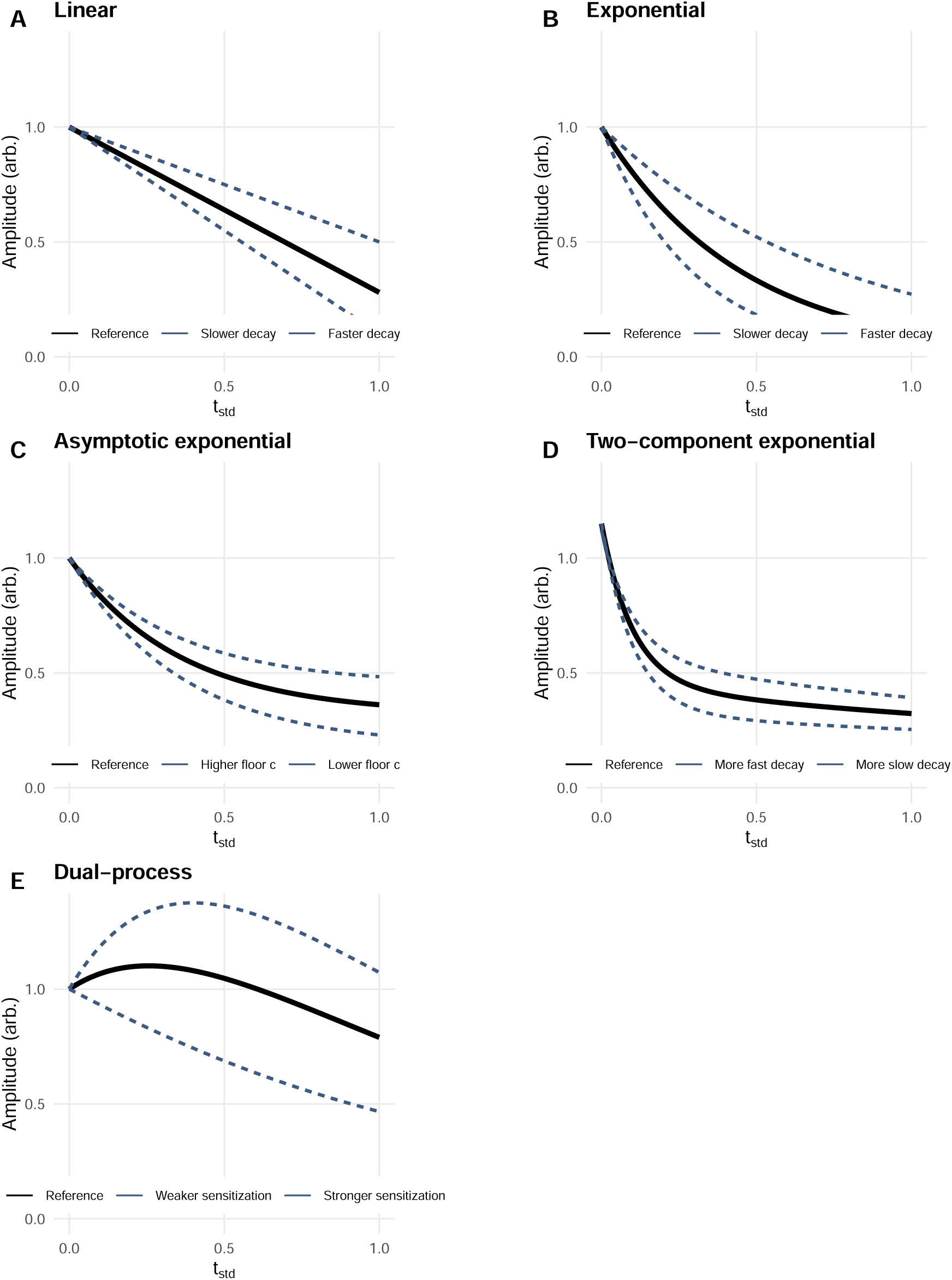
Schematic candidate habituation models on standardized session progress (*t* = *t*_std_ ∈ [0, 1]). Panels A–E: linear, exponential, asymptotic exponential, two-component expo-nential, and dual-process (2×3 grid, equal panel size). Black curves are reference parameters; blue curves change one parameter each. Amplitude is in arbitrary units on a shared scale.

**Table 2:** Candidate habituation curve models.

| Model | Functional form |
| --- | --- |
| Linear | $y = a + b t$ |
| Exponential | $y = a e^{-b t}$ |
| Asymptotic exponential | $y = c + a e^{-b t}$ |
| Two-component exponential | $y = c + a_1 e^{-b_1 t} + a_2 e^{-b_2 t}$ |
| Dual-process | $y = c + h e^{-b_h t} + s t e^{-b_s t}$ |

The five families differ in what shape they represent and in how many free parameters *p* they spend doing so. A linear fit (*p* = 2) describes a roughly constant decrement across the session; because it is unbounded below, we treat it as a descriptive benchmark rather than a process model, noting that the updated characteristics of habituation list linear as well as exponential decrements (Rankin et al., 2009). A simple exponential (*p* = 2) describes proportional loss of responding but forces the curve toward zero. An asymptotic exponential (*p* = 3) adds a floor *c*, so that responding need not approach zero^1^. A two-component exponential (*p* = 5) sums a fast and a slow decay and can capture a long tail without an early rise. The dual-process form (*p* = 5) is a *parametric* embodiment of Groves–Thompson ideas (Groves and Thompson, 1970): a decaying habituation term (*he*^−^*^bhx^*) together with a transient, sensitization-like term (*sxe*^−^*^bsx^*) that can produce an early rise before decline. These readings describe what a *winning* fit would mean if taken literally; the Results ask the narrower question of whether the same family describes the group average and the plurality of individuals.

### Model fitting and selection

All fitting was carried out on retained trials only. The linear model was fit by ordinary least squares; the four nonlinear families were fit by Levenberg–Marquardt nonlinear least squares (nlsLM in the R package minpack.lm). Because nonlinear fits are sensitive to where the optimizer starts—an under-converged fit can make a flexible family appear to fit worse than a simpler one—each nonlinear family was fit from a battery of starting values: three data-driven sets derived from the observed minimum, maximum, mean, and terminal values of the series, several randomized perturbations of those sets, and a final refinement started from the best solution found. Each attempt allowed up to 1,000 iterations under nonnegative lower bounds on all rate and amplitude parameters, and the solution with the lowest residual sum of squares across all attempts was retained for that family. A series was fit only if it contained at least eight retained trials and at least four distinct values of *t*_std_; shorter series were left unmodeled.

Among the five candidates, the one with the lowest small-sample corrected Akaike Information Criterion (AICc) was recorded as the *winner* for that series (Hurvich and Tsai, 1989; Burnham and Anderson, 2002). AICc balances fit against complexity: a more flexible curve is not preferred unless it lowers residual error enough to justify its additional parameters.

For each series,

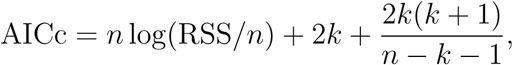

where *n* is the number of retained trials, RSS is the residual sum of squares, and *k* = *p* + 1 counts the *p* free curve parameters plus the residual variance (*k* = 3 for the linear and exponential families, 4 for the asymptotic exponential, and 6 for the two-component and dual-process families). Ties in AICc were broken alphabetically by model name. The rule is deliberately descriptive: it identifies which candidate best fits each series under a fixed penalty for the number of free parameters, not which family generated the data.

### Robustness and sensitivity analyses

Five auxiliary analyses asked whether the central mismatch could be an artifact of the analysis rather than a property of aggregation. The first concerns missing trials, which might make an exponential decay look linear when only the steep early portion of a session is observed (Appendix C). The second concerns the winner rule, which forces each subject into a single label even when two families fit almost equally well; a soft mixture analysis estimates population shares for each family without that hard assignment (Appendix D). The third concerns the small Study 3 cells (*n* = 20), where bootstrap resampling tested whether the plurality labels were stable (Appendix E). The fourth refit every series after excluding trials with |*z*| *>* 2 within trial number, to check that the pattern did not depend on extreme single-trial amplitudes. The fifth is a Monte Carlo simulation asking whether averaging a mixture of simple individual trajectories can by itself produce a more complex group-average curve. All five are reported together under the matching heading in the Results.

### Primary analysis: within-experiment group averages

The primary comparison was made within each of the four Experiment × ITI cells in Table 1, without mixing experiments. For every subject, all five families were fit to that subject’s retained trials and the winner was recorded; the *modal individual winner* for a cell is the family that won most often across its subjects—a plurality, not necessarily a majority. For the same cell, response amplitude was then averaged across subjects at each trial number, an unweighted mean over the participants with a retained value at that trial, and the five families were fit to the resulting group-average curve. We say a cell shows a *match*when the average-curve winner and the modal individual winner belong to the same functional family, and a mismatch otherwise. This within-cell comparison is the main basis for inference: it asks only that the same fitting rules be applied in parallel, not that the experiments share ITI, trial count, or stimulus.

### Secondary analysis: cross-experiment pooling

To ask whether aggregation distortions grow when experiments are combined, we formed pooled average curves across all 157 participants under six predefined schemes, crossing three ITI scopes (all cells; short ITI, 8 and 10 s; long ITI, 30 s) with two weighting rules. The rules answer different questions. Subject-weighted pooling estimates the average curve for the pooled set of participants, so that larger cells contribute more simply because they contain more subjects. Cell-equal-weight pooling instead gives each Experiment × ITI cell the same influence before fitting, which isolates whether any pooled result depends on the uneven cell sizes (20 to 67 participants).

Because the studies differ in session length (50 vs. 100 trials), pooling required a common axis: each curve was linearly interpolated onto 50 equally spaced points on [0, 1] for *t*_std_ (approx in R, with endpoint extrapolation) before averaging. Let *y_ig_*(*t*) denote the interpolated response for subject *i* in cell *g*, which has *n_g_* subjects. For a pooled set of cells G, subject-weighted pooling is

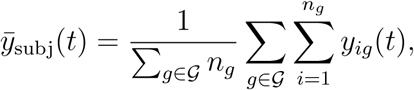

so that every subject contributes one curve and larger cells contribute more of them; in the short-ITI pool, for instance, Study 2 (*n* = 67) weighs more heavily than Study 3 at 10 s (*n* = 20). Cell-equal-weight pooling first averages within each cell and then averages those cell means,

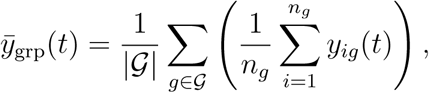

so that the two short-ITI cells count equally despite their different sizes. The same five families and selection rule were applied to each pooled curve.

### Transparency and openness

This study was not preregistered. All data and analysis code are publicly available in the project repository (https://github.com/omadav/habituation_curves); a Zenodo archive with a permanent DOI will be released at acceptance. All procedures were approved by the Ethics Committee of Universidad de Talca, and all participants provided written informed consent prior to participation. Large-language-model assistance (Claude Code, Anthropic) was used in code development, grammar checking and polishing for non-native authors; all analyses, interpretations, and conclusions are the authors’ own and were verified manually.

## Results

### Descriptive results

All four LED cells showed a clear average decrement across the session. However, individual subjects differed substantially in starting amplitude and rate of decline, which are the conditions under which aggregation artifacts are expected (Estes, 1956). Across all 157 participants, 46 (29%) were best described by linear decay, 44 (28%) by simple exponential, 38 (24%) by dual-process, 27 (17%) by asymptotic exponential, and 2 (1%) by two-component exponential. No single model commanded a majority in any cell. In Study 3 (30-s ITI) the plurality was dual-process (7 of 20 subjects, 35%), tied with exponential (7) and ahead of linear (5) and two-component (1). Figure 5 shows the full distribution of individual winning models within each of the four cells, and Figure 3 plots the underlying individual trajectories against the group average.

**Figure 3:**
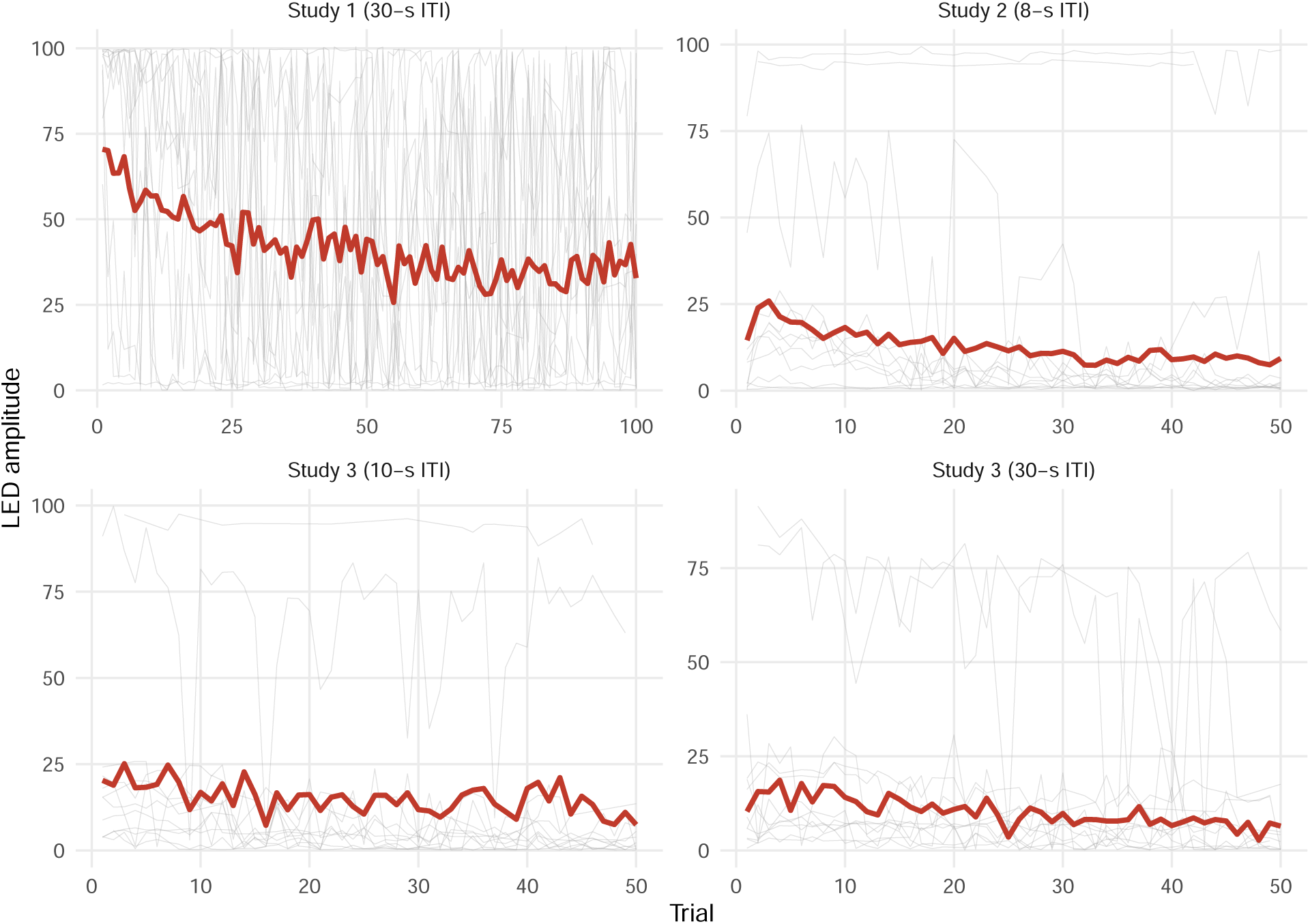
A random sample of up to 12 individual LED trajectories (grey) together with the group-average curve over all participants (red), for each Experiment × ITI cell. The individual curves span a wide range of starting amplitudes and decline rates, so that the average traces the central tendency of a heterogeneous set rather than a representative individual trajectory.

### Individual-level vs. average-curve model comparison

Table 3 summarizes the comparison for each LED cell. The modal individual winner was linear in three of the four cells (Studies 1, 2, and 3 at 10-s ITI), capturing 28% to 35% of subjects. In Study 3 (30-s ITI) the plurality was dual-process (7 of 20 subjects, 35%), tied with exponential (7) and ahead of linear (5).

**Table 3:** Individual-level and average-curve model comparison for the LED Experiment × ITI cells.

| Cell | $N$ | Modal individual | $n$ (%) | Average-curve model | Match |
| --- | --- | --- | --- | --- | --- |
| Study 1, 30-s ITI | 50 | Linear | 14 (28%) | Asym. Exp. | No |
| Study 2, 8-s ITI | 67 | Linear | 20 (30%) | Dual Process | No |
| Study 3, 10-s ITI | 20 | Linear | 7 (35%) | Exponential | No |
| Study 3, 30-s ITI | 20 | Dual Process | 7 (35%) | Exponential | No |

Modal individual = most frequent best-fitting model (plurality). Match = same functional family for average and modal winners. Free curve parameters: linear and exponential, $p = 2$ ; asymptotic exponential, $p = 3$ ; two-component exponential and dual-process, $p = 5$ . In Study 3 (30-s ITI) dual-process and exponential were tied among individuals (7 each); ties are broken alphabetically.

By contrast, the best-fitting model for the group average was asymptotic exponential in Study 1, dual-process in Study 2, and exponential in both Study 3 cells. In every cell this differed from the modal individual winner—a **shape mismatch**, a different functional family than the plurality, in all four cells. In Studies 1 and 2 the mismatch was also a **parameter-count mismatch**: the average selected a model with strictly more free parameters than the modal individual (*p* ≥ 3 vs. *p* = 2). Study 3 (10-s ITI) showed shape mismatch without higher parameter count—linear and exponential both have *p* = 2, but the average curved exponentially whereas the plurality declined approximately linearly. In Study 3 (30-s ITI) the average selected exponential while the individual plurality was dual-process (35%; 95% CI 15–55, Table 9). No cell’s average matched its individual plurality.

Figure 4 illustrates the contrast across cells. Study 1 (Panel A) is the clearest example: the individual plurality is linear, but the group average curves toward an asymptote.

**Figure 4:**
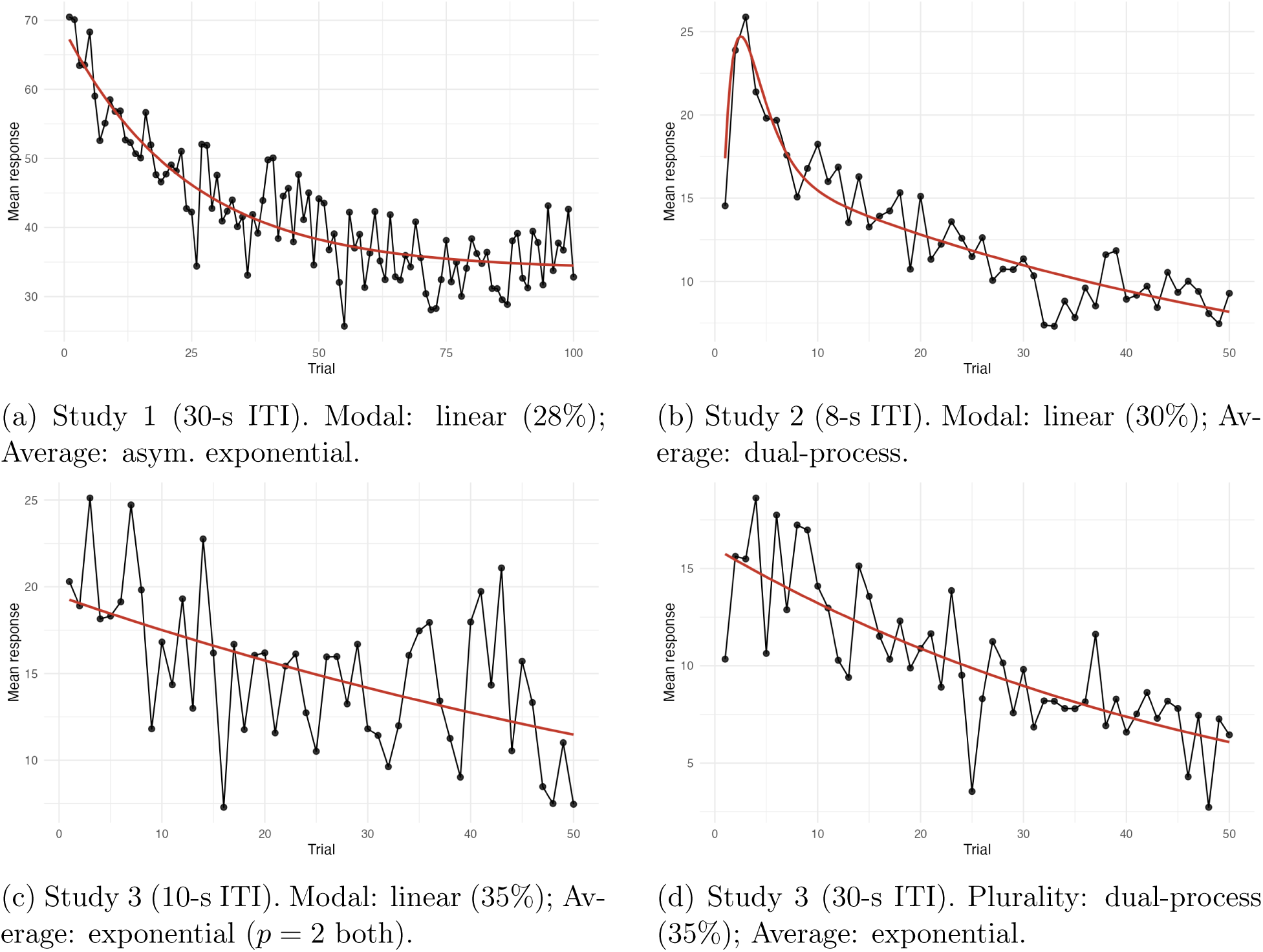
Group-average curves with best-fitting model for each Experiment × ITI cell.

**Figure 5:**
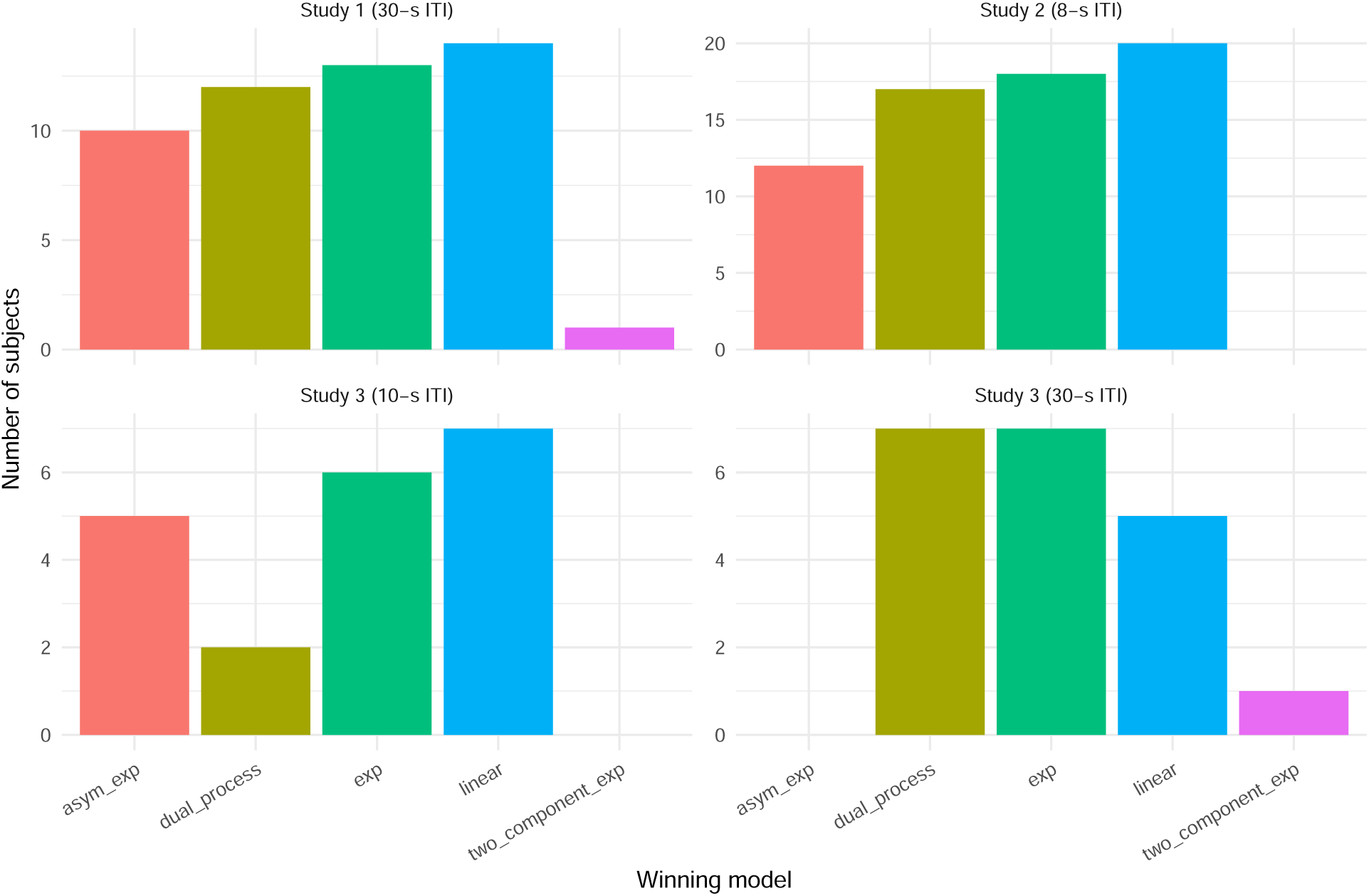
Distribution of individual winning models across the four LED Experiment × ITI cells (*N* = 157).

### Pooled analyses

Pooling across cells amplified the discrepancy. The subject-weighted analysis asked what happens when every participant is allowed to contribute equally to the pooled curve; under that rule, larger cells necessarily have more influence. In the all-group and short-ITI subject-weighted pools the modal individual winner was linear; in the long-ITI pool it was exponential. In every subject-weighted pool the pooled average was nonetheless best described by dual-process. The cell-equal-weight analysis asked a different question: whether the same conclusion holds when each Experiment × ITI cell contributes equally, regardless of sample size. This matters because the cells were uneven, ranging from 20 to 67 participants. If the pooled dual-process result were driven only by the largest cell, it should weaken or disappear when each cell is given the same weight. Instead, the full-sample and short-ITI equal-weight pools again selected dual-process. The long-ITI equal-weight pool selected asymptotic exponential, but it too mismatched the modal individual winner (exponential). In no pooled comparison did the average-curve winner match the modal individual winner.

Dual-process was the best descriptor of most pooled averages, yet it was the modal individual family in none of these pools (Table 4).

**Table 4:** Pooled LED analyses: modal individual winner vs. pooled average-curve winner.

| Pool | $N$ | Cells | Modal individual | Average winner |
| --- | --- | --- | --- | --- |
| All, subject-weighted | 157 | 4 | Linear | Dual Process |
| Short-ITI, subject-weighted | 87 | 2 | Linear | Dual Process |
| Long-ITI, subject-weighted | 70 | 2 | Exponential | Dual Process |
| All, equal-weight | 157 | 4 | Linear | Dual Process |
| Short-ITI, equal-weight | 87 | 2 | Linear | Dual Process |
| Long-ITI, equal-weight | 70 | 2 | Exponential | Asym. Exp. |

Subject-weighted: equal subject weight. Equal-weight: equal Experiment × ITI cell weight before averaging.

### Robustness and sensitivity analyses

Together, these analyses supported the main conclusions. Missing trials showed only a weak, non-significant association with linear classification, and simulations indicated that missingness at the observed levels could not account for the observed linear plurality. A soft-classification approach produced the same qualitative mismatch in the main cells, and bootstrap resampling showed that modal shares were uncertain, especially in the smaller Study 3 cells. Refitting after excluding within-trial outliers (|*z*| *>* 2) left the pattern unchanged. Full details of the missing-trial, soft-classification, and bootstrap checks are reported in Appendices C, D, and E, respectively; the fifth analysis, a generative simulation, is described next.

A separate question is mechanistic rather than defensive: could the complex group averages arise from aggregation alone, even when most individuals are simpler? The two most common individual families were linear (29%) and simple exponential (28%), yet the pooled average selected dual-process. We therefore asked whether mixing only those two simple families is enough to push the average into a more complex family. For each replicate (*R* = 300 per condition) we generated a synthetic group of *N* = 80 subjects, each drawn from a linear (*y* = *a* + *bt*) or a simple exponential (*y* = *ae*^−^*^bt^*) generator with parameters drawn from fixed normal distributions chosen to span a realistic range of starting amplitudes and decay rates; the proportion of linear subjects *p*_lin_ was varied from 0 to 1 in steps of 0.1, with the remainder exponential. The group average of each synthetic sample was fit with the four candidate families (linear, exponential, asymptotic exponential, and dual-process) under the same selection rule used throughout. No asymptotic-exponential or dual-process subjects were ever included, so any selection of those families at the group level is produced by aggregation rather than by individuals (Table 5).

**Table 5:** Modal winning model for the group average as a function of the proportion of linear subjects in a mixture of linear and exponential individuals (*N* = 80, *R* = 300 replicates per row). No asymptotic exponential or dual-process individuals were included.

| $p_{\text{lin}}$ | Linear (%) | Exp (%) | Asym-exp (%) | Dual (%) | Modal winner |
| --- | --- | --- | --- | --- | --- |
| 0.0 | 0.0 | 0.0 | 83.0 | 17.0 | Asym-exp |
| 0.1 | 0.0 | 0.0 | 92.7 | 7.3 | Asym-exp |
| 0.2 | 0.0 | 0.3 | 91.3 | 8.3 | Asym-exp |
| 0.3 | 0.0 | 1.0 | 90.7 | 8.3 | Asym-exp |
| 0.4 | 0.0 | 9.0 | 82.7 | 8.3 | Asym-exp |
| 0.5 | 0.0 | 41.3 | 53.7 | 5.0 | Asym-exp |
| 0.6 | 0.0 | 87.3 | 10.0 | 2.7 | Exponential |
| 0.7 | 0.0 | 89.3 | 0.0 | 10.7 | Exponential |
| 0.8 | 0.3 | 40.0 | 0.0 | 59.7 | Dual-process |
| 0.9 | 32.0 | 6.3 | 0.0 | 61.7 | Dual-process |
| 1.0 | 92.3 | 0.0 | 0.0 | 7.7 | Linear |

The pattern is clear (Table 5). When the group is entirely exponential (*p*_lin_ = 0), the average is best described by an asymptotic exponential, because exponentials with heterogeneous rates average into a slower decline with an apparent floor (a consequence of Jensen’s inequality; Appendix F). As *p*_lin_ rises to 0.6–0.7 the average reverts to a simple exponential, and at *p*_lin_ = 0.8–0.9 it selects a dual-process model in about 60–62% of replicates even though no individual is dual-process: a mixture of linear decline and residual exponential curvature produces an average with an early exponential phase and a later near-linear tail, which is exactly what the dual-process form captures.

The empirical pooled composition (29% linear, 28% exponential) sits near the region where the average selects asymptotic or simple exponential, consistent with what the pooled curve returned; the asymptotic-exponential and dual-process individuals plausibly carry it the rest of the way to dual-process. The point is not that dual-process individuals are absent—about a quarter of participants are best fit by that family—but that the group average over-selects it: the pooled curve reads as dual-process in every cell, whereas dual-process is the individual plurality in only one. Family-level heterogeneity is sufficient to manufacture a dual-process group average, so the group-average shape cannot be read as the shape of the typical individual. The simulation addresses the direction of mismatch observed in most cells and all pooled analyses, where the average is more complex than the individual modal. Study 3 (30-s ITI) shows the reverse: the average selected a simpler family (exponential) than the individual modal (dual-process, tied with exponential at 7 subjects each), which falls outside the scope of this particular demonstration but underscores the same general point—the average can diverge from the individual distribution in either direction.

## Discussion

In this study, using habituation data from our research group, we set out to investigate if average curves represent the plurality of individual fits. The main finding is that the group-average winner disagreed with the individual-level plurality in all four cells. In Studies 1, 2, and 3 (10-s ITI), the modal individual was linear, but the average selected asymptotic exponential, dual-process, or simple exponential, respectively. Study 3 (10-s ITI) is notable because both families have the same parameter count (*p* = 2); the average curved where most individuals declined approximately linearly. Study 3 (30-s ITI) was the only cell whose individual plurality was itself dual-process, yet its average selected exponential—so the two summaries disagreed even there. Pooling made the discrepancy worse: every pooled average was best described by dual-process or asymptotic exponential, while the overall modal individual was linear (29%). Thus, the central result is not only that group averages can select a different family than the most frequent individual family, but also that no single functional form characterized most participants. The individual-level distribution itself is therefore part of the finding: in this preparation, the “typical” habituation curve was not well represented by a single dominant curve family.

This pattern is consistent with the aggregation argument made by Estes (1956) and formalized by Ashby et al. (1994): when individuals differ in a rate parameter, the group average can have a different functional form than any individual curve. For exponential decays, heterogeneous rates produce an average that drops quickly at first and then flattens, because fast decayers pull the early mean down while slow decayers keep the late mean elevated — a consequence of Jensen’s inequality (Appendix F). By contrast, the mean of linear functions is linear, so a group average that selects a more complex model implies that at least some individuals are nonlinear. The simulation confirmed this directly. This does not imply that dual-process dynamics are absent at the individual level; it implies that a dual-process group average cannot, by itself, establish that they are present.

The robustness and sensitivity analyses told a consistent story: each alternative explanation contributed something, but none removed the mismatch. Missing trials were only weakly and non-significantly associated with linear classification (*p* = .072), and dropout at observed severity could not reproduce the 29% linear rate (Appendix C). Allowing fractional membership across curve families did not make dual-process the dominant individual process in the cells where the average selected it (Study 2: 30% soft dual-process against an exponential-led soft estimate of 52%, despite a dual-process average; Appendix D). Bootstrap resampling confirmed that the plurality labels, though uncertain in the small Study 3 cells, never reached a majority for any of the more complex families (Appendix E). The mismatch therefore does not appear to depend on missing trials, on forcing a single label per subject, or on sampling noise in the smaller cells.

These results have direct implications for how dual-process accounts of habituation are evaluated from response curves. A curve that rises early and then declines has often been taken as evidence that two opposing processes are operating: a decremental habituation process and an incremental sensitization process (Groves and Thompson, 1970). The present findings do not challenge the existence of sensitization, nor do they imply that repeated stimulation cannot produce both response-decremental and response-potentiating effects under appropriate conditions. They show instead that a dual-process-shaped group curve is not, by itself, sufficient evidence that the typical individual trajectory was generated by two opposing processes. When individual trajectories are heterogeneous, aggregation can produce an average curve with an apparent dual-process form even when simpler individual trajectories are common. Thus, group-average curve shape should be used cautiously as evidence for dual-process dynamics unless it is accompanied by individual-level analyses showing that the same form is present in a substantial proportion of participants.

The preparation analyzed here is also relevant for interpreting this point. The data come from human eyeblink responses elicited by repeated stimulation in a relatively constrained startle-habituation setting. Such a preparation may be less likely to produce strong or sustained sensitization than procedures using more intense, aversive, or motivationally significant stimuli. Thus, the absence of a dominant dual-process individual profile in these data should not be generalized to all forms of habituation or all response systems. It is entirely possible that stronger sensitization-like increments would be observed in other preparations, species, stimulus intensities, or emotional contexts. The present contribution is methodological: it shows that the group-average curve can overstate the prevalence of a complex curve shape among individuals.

The findings also bear on the long-standing distinction between acquisition curves and retention tests. Several authors have argued that habituation, as a learning phenomenon, is best evaluated through retention, common-test, specificity, or dishabituation procedures because within-session curves can confound learning with performance (e.g., Colwill et al., 2023). The present analysis does not weaken that concern. It instead addresses a complementary question: when researchers do analyze the habituation curve as a description of response change during repeated stimulation, should the group-average curve be treated as representative of individual curve shape? Our results suggest that it should not. At the same time, the short-lived dynamics expressed during acquisition should not be dismissed as theoretically unimportant. Transient response decrement may reflect short-term memory for recent stimulation, priming, or other stimulus-processing mechanisms, even when it does not provide a pure measure of durable learning. This point is consistent with contemporary analyses of habituation theories in which within-session decrement and retention can depend on separable short- and long-term mechanisms (Jorquera et al., 2025; Becerra et al., 2026). The key issue, which was outside the scope of the present article, is to determine when such short-term changes reflect memory-like processes and when they can be explained by sensory adaptation, motor fatigue, or other nonmnemonic factors.

This distinction is especially important for clinical and individual-difference research. Patient-control studies often use habituation curves to characterize phenotypes, but clinical samples are rarely homogeneous. In anxiety and stress-related disorders, clinical differences in habituation may appear in initial response level, rate of change, total decrement, or retention, and these components do not necessarily covary (Cancino et al., 2026). The present results add a further warning: even before comparing groups, the curve fitted to the average may fail to represent the distribution of individual trajectories within a group. In clinical contexts, such heterogeneity should not be treated only as noise. It may be part of the phenomenon to be explained, especially when different individuals express different combinations of initial responsiveness, decrement rate, asymptote, response increases, or delayed recovery.

Several limitations should be noted. The analysis combined studies that differed in ITI, trial count, and stimulus; primary inferences rest on within-cell comparisons, and pooling required interpolation across session lengths. Study 3 cells contained only 20 participants, and bootstrap intervals confirm that modal labels are unstable plurality descriptors in those cells (Appendix E). The candidate set was fixed and descriptive; agreement or disagreement between winners does not establish that any parametric family is the true generative process. In addition, the analysis was restricted to the first uninterrupted repeated-stimulation phase of each dataset and did not include independent tests of retention, stimulus specificity, or dishabituation. The present findings therefore concern curve-shape inference during acquisition, not the full set of criteria needed to establish habituation as learning.

The conclusions are also bounded by the preparation analyzed here. All analyses were based on human eyeblink responses in a relatively constrained startle-habituation procedure using a mild eliciting stimulus. This setting may be less likely to generate strong or sustained sensitization than procedures involving more intense, aversive, or motivationally significant stimulation. Accordingly, the absence of a dominant dual-process individual profile in these data should not be generalized to all forms of habituation. Stronger sensitization-like increments may well be observed in other preparations, species, response systems, stimulus intensities, or emotional contexts. The methodological point remains that, whenever such curve forms are evaluated, the group-average curve can overstate the prevalence of a complex trajectory among individuals.

### Recommendations for reporting habituation curves

The following recommendations extend those of McDiarmid et al. (2017) to within-session curve-shape inference. The concern is practical as well as methodological: clinical studies comparing habituation across patient and control groups (Cancino et al., 2026) face the same problem when group-average fits are interpreted as descriptions of individual processes.

1. Fit candidate models to individual trial series, not only to the group mean.
2. Report the distribution of winning models (counts or proportions for every candidate family), not only the modal winner.
3. Fit the same model set to the group-average curve and state explicitly whether its winner matches the individual plurality. In group-comparison studies (e.g., clinical vs. control groups), also examine whether differences in average curve shape reflect differences in individual trajectories, differences in the mixture of curve types, or both.
4. Use simulation and sensitivity analyses (e.g., missingness checks, soft mixture estimates) to test whether aggregation or dropout could bias shape inferences before interpreting the average curve as a description of the typical participant.

These recommendations are not equivalent to adding random effects to a trajectory model. Random effects are useful when the analyst has already specified a common functional form and wishes to estimate how its parameters vary across participants. The present issue concerns an earlier step: whether a common functional form adequately describes the individuals in the first place. A random intercept or random slope can capture differences in initial level or rate of change, and nonlinear random effects can capture variation in parameters such as decay rate or asymptote, but all participants remain constrained by the same curve family. By contrast, the present approach asks whether different individuals are better described by different candidate families, and whether the family selected for the group average represents the distribution of individual fits. Mixed-effects models and individual-level model selection are therefore complementary: the former estimates parameter variability within a chosen form, whereas the latter evaluates heterogeneity in functional form itself.

To make these recommendations concrete, consider Study 1. Fitting the five families to each of its 50 participants returns a *distribution* of winners—linear for 28% of subjects, with the rest divided among the exponential, asymptotic-exponential, and dual-process families, and no family holding a majority. Fitting the same five families to the 50-subject average curve, by contrast, selects an asymptotic exponential. Reported side by side, the two summaries make the mismatch legible: the average curve is more complex than the most common individual, and the asymptotic shape should not be read as the trajectory of the typical participant. Reporting the average curve alone would have concealed that gap.

In sum, the present results suggest that habituation group averages can select curve families that differ from the most common individual form. This finding does not reduce the value of habituation curves; it clarifies the level at which they should be interpreted. The average curve provides a useful description of the ensemble, whereas the individual-level distribution indicates how response dynamics are organized across participants. Individual-level model comparison is feasible with standard nonlinear least squares, at least in datasets of this kind, and can be reported alongside the group-average fit. By combining these two levels of analysis, researchers can preserve the theoretical and empirical value of habituation curves while avoiding the assumption that the average curve represents the modal individual pattern.

## Data Availability

The analysis code, de-identified trial-level LED amplitudes, and derived outputs needed to reproduce all tables and figures are publicly available in the project repository: https://github.com/omadav/habituation_curves. A Zenodo archive with a permanent DOI will be released at acceptance; the repository URL is the access point at submission.

## Code Availability

All code will be released under a creative commons license at the same repository. The pipeline requires R ≥ 4.0 with packages tidyverse, minpack.lm, and readxl. The main individual-versus-average analysis—every per-cell winner, the pooled comparison, the bootstrap and soft-mixture checks, and the corresponding figures—is reproduced end to end by a single script, robust_analysis.R, which uses the robust fitting described in the Method.

## Supporting information

Translational Abstract

## Appendix A Archived Experiment Protocols

The main analysis pooled LED habituation series from three archived studies conducted in the Vogel laboratory. What follows is the protocol detail needed to interpret the four analysis groups in Table 1. Stimulus-specific and context manipulations were not entered into the curve-shape comparison; they are recorded here for completeness.

### Study 2

Participants were students tested on the Eyeblink Conditioning System (EBC). The session analyzed here was the first habituation block (50 trials). The intertrial interval was 8 s. On each trial, a 50 ms blink-eliciting event was delivered: white noise at 105 dB or an air puff at 0.1 MPa, counterbalanced across participants. LED eyelid closure and EMG were recorded; only LED is analyzed in the main text (EMG in Appendix B).

### Study 3

Participants were tested on the EBC with a 50 ms air puff at 0.2 MPa. Two habituation ITI conditions were run as separate cells: 10 s and 30 s, each with 50 trials in Session 1. The main text retains only the no-distractor Control arm, without dishabituation probes, test-phase trials, or Session 2 data, so each series is uninterrupted repeated habituation. LED and EMG were recorded.

### Study 1

Fifty archived LED records came from a context eyeblink study (100 habituation trials, 30 s ITI). Habituation stimuli were air puff or tone, crossed with same versus different context labels across phases of the study. Participant identifiers were assigned at harmonization because the legacy file did not provide a stable subject key. Only Session 1 habituation amplitudes entered the reanalysis.

### Shared recording and trial metric

Across studies, the infrared LED signal was sampled at 1 kHz, low-pass filtered at 40 Hz offline, and scored trial-wise from 500 ms before onset to 300 ms after onset. Baseline activity in the prestimulus window was subtracted; the dependent variable was the peak post-onset amplitude after baseline correction. The same rules were applied when building group-average curves.

## Appendix B EMG Replication

The LED analysis was repeated, with the same robust fitting, on the EMG measure available from Studies 2 and 3 (*N* = 106 across three Experiment × ITI cells). The average-curve winner was dual-process in all three cells. The modal individual model was dual-process for Study 2 (33% of 66 subjects) and simple exponential for both Study 3 cells (45% of 20 subjects each). The average therefore mismatched the individual plurality in the two Study 3 cells; in Study 2, both the plurality and the average were dual-process, the one EMG cell in which the two summaries agreed. The EMG data thus reproduce the LED pattern—an average more complex than, or different from, the individual plurality—in two of the three cells, in a different response modality. The exception is instructive rather than awkward. The average recovered dual-process exactly where the individual plurality was already dual-process; the group mean tracks the individual form when the population is homogeneous in that form, and departs from it only when the population is mixed. The mismatch documented throughout this paper is therefore a symptom of heterogeneity, not a property of averaging as such.

## Appendix C Missing-Trials Robustness Details

Individual-level model selection classified 29% of subjects as linear decayers. A natural concern is that this could be a measurement artifact: subjects with more missing late trials might appear linear because only the steep early portion of an exponential decline is observed. Linear winners did show slightly higher mean missingness (19.8%) than exponential winners (18.0%), but a logistic regression predicting linear classification from proportion missing, experiment, and ITI condition returned only a weak, non-significant association (*β* = 0.24 per 10 percentage points, *SE* = 0.13, *z* = 1.80, *p* = .072). To gauge how much false-linear classification missingness could produce, we simulated truly exponential subjects under independent dropout (MCAR), late-trial-biased dropout (MNAR-late), and hard truncation (Figure 6). At missingness levels typical of empirical linear winners (mean 19.8%), the most adversarial mechanism produced at most 11.0% linear misclassification, well below the observed 29%; only at an extreme 50% missing did any mechanism approach the empirical rate, and only hard truncation (31.0%; Table 7). Missingness therefore cannot account for the observed linear plurality.

**Figure 6:**
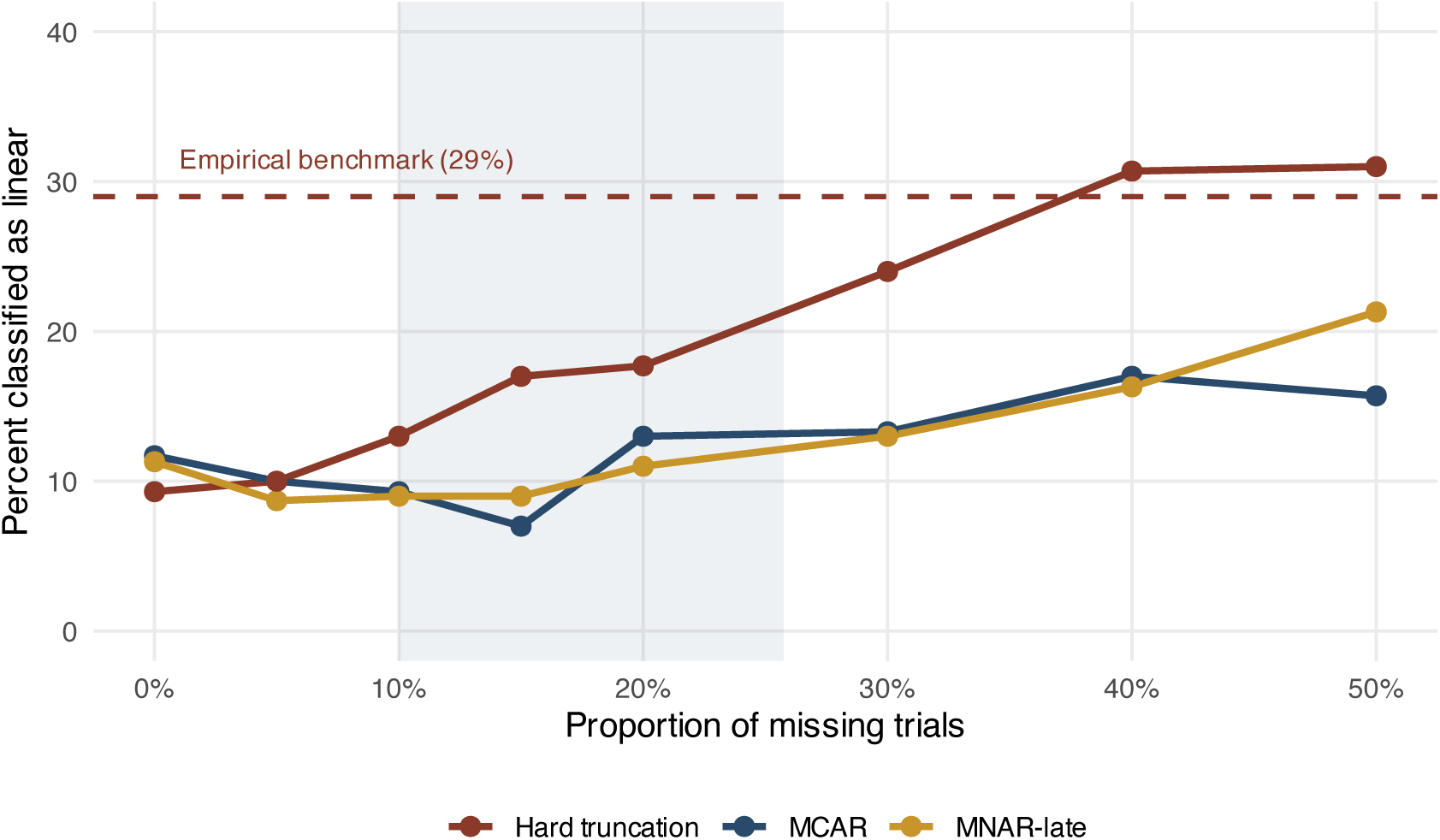
Proportion of truly exponential simulated subjects misclassified as linear as a function of *p*_miss_. Dashed line: empirical benchmark (29%). Shaded band: interquartile range of missingness among empirical linear winners (mean 19.8%).

**Figure 7:**
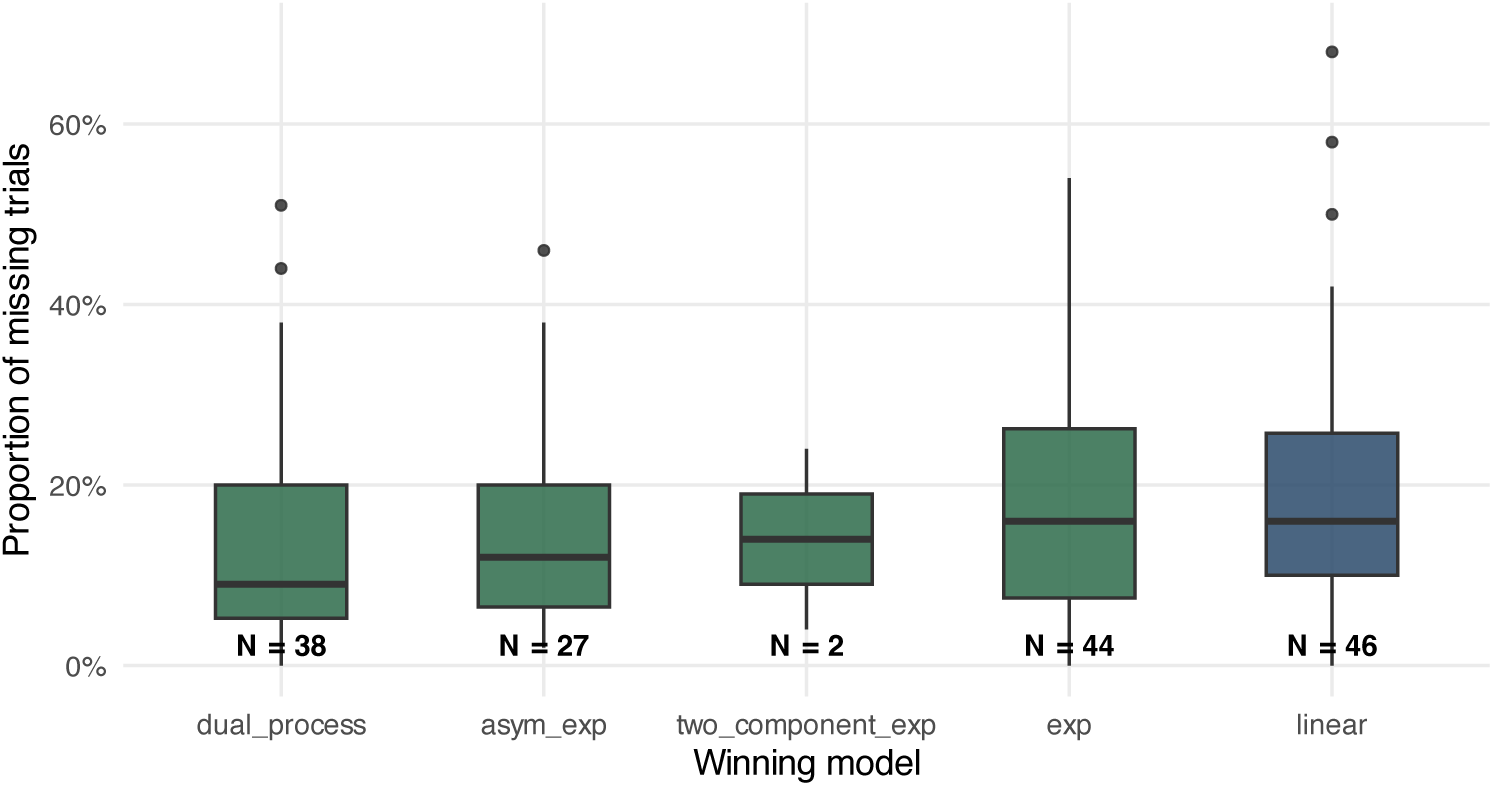
Proportion of missing trials by winning model. Box labels show *N* per class.

**Table 6:** Observed trials and proportion missing by individual winning model (LED, *N* = 157).

| Winning model | $N$ | Mean observed trials | Mean % missing |
| --- | --- | --- | --- |
| Linear | 46 | 52.3 | 19.8% |
| Exponential | 44 | 52.5 | 18.0% |
| Dual-process | 38 | 55.8 | 13.6% |
| Asymptotic exponential | 27 | 58.5 | 14.7% |
| Two-component exponential | 2 | 62.0 | 14.0% |

Study 1: up to 100 nominal trials; Studies 2 and 3: 50. Counts include trials excluded before fitting.

**Table 7:** Missingness required to reproduce the empirical linear rate under a purely exponential data-generating process.

| Mechanism | $p_{\text{miss}}$ to reach $\geq 29\%$ linear | Max linear % at 50% missing |
| --- | --- | --- |
| MCAR | $> 50\%$ | 17.0% |
| MNAR-late | $> 50\%$ | 21.3% |
| Hard truncation | 50% | 31.0% |

## Appendix D Soft Mixture Analysis (One Label vs. Population Shares)

### Why this check is needed

The primary analysis gives each subject one winning curve family—whichever candidate had the lowest AICc in that cell. That rule is simple to report, but it is strict. When two models fit nearly equally well, a small change in noise can flip the assigned label, and it may well be that the reported plurality (e.g., 31% linear in Study 2) is unstable.

One possible way to test this potential brittleness of the results is relaxing the one-label-per-subject rule. If we allow each subject to belong to multiple curve families with different probabilities, does the *population* in a cell look like the family the group average selected? If these soft estimates showed, for example, that most individuals in Study 2 were dual-process, then a dual-process group average would not contradict the individual-level picture. The analysis below reweights the same model-comparison scores already obtained in the primary analysis, treating each family as a latent class whose relative support is estimated within each Experiment × ITI cell.

### Method

For each subject and each of the four primary families (linear, exponential, asymptotic exponential, dual-process), we converted the fitted model’s AICc to a score *ℓ_ik_* = −AICc*_ik_/*2 (higher is better). The two-component exponential was excluded because it was selected by fewer than 2% of subjects and was often unstable at *n* = 20. An expectation–maximization (EM) algorithm then estimated population shares *π*^*_k_* within each cell: the E-step computed each subject’s posterior probability of belonging to family *k*, and the M-step updated *π*^*_k_* as the average posterior. Iterations continued until the change in observed log-likelihood fell below 10^−8^. Because the scores carry the same complexity penalty as the primary analysis, a flexible family cannot dominate merely by having more parameters. The few subjects with a convergence failure in a given family received a very low score for that family.

### Results

Table 8 reports the estimated population *π*^*_k_* alongside the hard plurality and the shares average-curve winner for each cell.

**Table 8:** Soft mixture estimates: population share *π*^*_k_*(%) by curve family, compared with the hard plurality and the average-curve winner.

| Cell | EM $\hat{\pi}_k$ (%) | | | | Hard plurality | Average curve |
| --- | --- | --- | --- | --- | --- | --- |
|  | Lin. | Exp. | Asym. | Dual |  |  |
| Study 1, 30-s ITI | 24 | 33 | 14 | 30 | Linear (28%) | Asymptotic exp. |
| Study 2, 8-s ITI | 6 | 52 | 12 | 30 | Linear (30%) | Dual-process |
| Study 3, 10-s ITI | 0 | 63 | 30 | 7 | Linear (35%) | Exponential |
| Study 3, 30-s ITI | 29 | 27 | 0 | 44 | Dual-process (35%) | Exponential |
| Pooled ($N = 157$) | 14 | 44 | 12 | 29 | Linear (29%) | Dual-process |

EM proportions estimated separately within each cell; pooled row averages over all 157 subjects. Study 3 cells ( $n = 20$ ) are the most variable; zero entries reflect $\hat{\pi}_k$ converging to a boundary.

In Study 1 the hard plurality was linear (28%) but the soft estimate placed exponential and dual-process highest (33% and 30%); neither matched the average’s asymptotic exponential. In Study 2 the hard plurality was linear (30%) and the soft estimate exponential (52%), yet dual-process drew only 30% soft support despite a dual-process average. In Study 3 (10-s ITI) soft classification put exponential first (63%) while the hard plurality was linear, and the average was exponential. In Study 3 (30-s ITI) both the hard plurality and the soft estimate favored dual-process (35% and 44%) while the average selected exponential. Per-subject agreement between the hard-winner method reported in the main text and the soft modal assignment ranged from 55% (Study 3, 10-s ITI) to 90% (Study 3, 30-s ITI); the median maximum posterior was 0.69.

### Conclusion

The soft classification analysis rules out a specific artifact: the average-individual mismatch is not explained by forcing one label per subject while a hidden majority actually matches the average winner. In no cell does the soft estimate place the average’s family at a majority, so allowing fractional membership does not reconcile the individual and average summaries. The hard-winner method reported in the main text is therefore a reasonable summary of the individual-level composition.

## Appendix E Bootstrap Stability of Modal Shares

Because the Study 3 cells contain only *n* = 20 participants, we used a nonparametric bootstrap to ask how stable the modal (plurality) share is to which participants happen to be sampled. Within each Experiment × ITI cell we resampled participants with replacement (*B* = 5,000 resamples), keeping each resampled participant’s winning model fixed at the value assigned in the primary analysis; models were *not* refit. For each resample we re-computed the percentage of participants in every curve family. The reported 95% intervals are percentile intervals (2.5th–97.5th) of those bootstrap distributions; they describe the sampling variability of the share itself, not uncertainty in any individual fit. Table 9 gives the interval for the modal model and for the linear family in each cell. The intervals are wide, as expected at *n* = 20, but in no cell does the interval for asymptotic exponential or dual-process reach a majority, and in every cell the modal share is consistent with the plurality being a minority of the participants.

**Table 9:** Bootstrap 95% intervals for modal model share by Experiment × ITI cell (*B* = 5,000 resamples).

| Cell | $N$ | Modal model | Observed share | 95% CI | Linear share [95% CI] |
| --- | --- | --- | --- | --- | --- |
| Study 1, 30-s ITI | 50 | Linear | 28% (14/50) | 16–42 | 28% (16–42) |
| Study 2, 8-s ITI | 67 | Linear | 30% (20/67) | 19–42 | 30% (19–42) |
| Study 3, 10-s ITI | 20 | Linear | 35% (7/20) | 15–55 | 35% (15–55) |
| Study 3, 30-s ITI | 20 | Dual-process | 35% (7/20) | 15–55 | 25% (5–45) |

Percentile intervals from subject-level bootstrap resampling with replacement ( $B = 5,000$ ); duplicate draws retain full subject records.

## Appendix F Jensen’s Inequality and Average-Curve Curvature

Suppose a subset of subjects habituate as simple exponentials with heterogeneous rate pa-rameters,

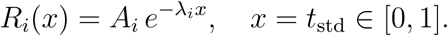

The group-average curve at each point is

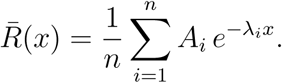

For fixed *x >* 0 and fixed *A*, the mapping *g*(*λ*) = *Ae*^−^*^λx^* has *g*^′′^(*λ*) = *Ax*^2^*e*^−^*^λx^ >* 0, so it is strictly convex. Jensen’s inequality therefore gives

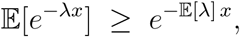

with equality only if all *λ_i_* are identical. Heterogeneous rates produce an average that decays more slowly than the exponential defined by the mean rate: fast decayers pull the early average down, but slow decayers keep the later average elevated, creating a flatter tail. In least-squares competition, this slower tail is well captured by an asymptotic exponential *y* = *c* + *ae*^−^*^bx^* (*c >* 0), explaining why the group-average curve selects that family even when few individuals show asymptotic dynamics.

The same logic generalizes beyond the exponential subset. The group-average curve is the superposition of all individual trajectories. If subjects include a mixture of linear, exponential, and asymptotic exponential decayers, the mean inherits contributions from each, and the combined trajectory can exhibit curvature that a more complex model (asymptotic exponential, dual-process) captures well. By contrast, if all subjects were truly linear, *R_i_*(*x*) = *a_i_* + *b_i_x*, then *R̄*(*x*) = *ā* + *̄b x* remains linear, because the mean of lines is a line. The appearance of nonlinear curvature in the group mean therefore indicates the presence of at least some nonlinear individual trajectories, but does not imply that the *typical* individual is asymptotic or dual-process.

## Footnotes

1 We do not commit to an explanation of this feature. The floor may reflect a genuine biological constraint on responding, or it may be a measurement artifact of the apparatus.

## References

Ashby, F. G., Maddox, W. T., and Lee, W. W. (1994). On the dangers of averaging across subjects when using regression or covariance models to interpret group data. Journal of Experimental Psychology: General, 123(2):168–175.

Becerra, S. A., Pinto, J. A., Jorquera, O. E., Matamala, P. D., Ramírez, C. C., and Vogel, E. H. (2026). Interstimulus interval effects on habituation: A systematic review with theoretical implications. Psychological Bulletin, 152(1):96.

Burnham, K. P. and Anderson, D. R. (2002). Model Selection and Multimodel Inference: A Practical Information-Theoretic Approach. Springer, New York, NY, 2 edition.

Cancino, N. A., Farfán, O. M., Campos, S. V., Ramos, N. A., Kreither, J., and Vogel, E. H. (2026). What do patient–control studies of habituation in anxiety and stress disorders reveal? Psychophysiology, 63(5):e70315.

Colwill, R. M., Lattal, K. M., Whitlow Jr., J. W., and Delamater, A. R. (2023). Habituation: It’s not what you think it is. Behavioural Processes, 207:104845.

Davis, M. and Wagner, A. R. (1968). Startle responsiveness after habituation to different intensities of tone. Psychonomic Science, 12(7):337–338.

Estes, W. K. (1956). The problem of inference from curves based on group data. Psychological Bulletin, 53(2):134–140.

Farrell, S. and Lewandowsky, S. (2018). Computational Modeling of Cognition and Behavior. Cambridge University Press, Cambridge, UK.

Gallistel, C. R., Fairhurst, S., and Balsam, P. (2004). The learning curve: Implications of a quantitative analysis. Proceedings of the National Academy of Sciences, 101(36):13124–13131.

Groves, P. M. and Thompson, R. F. (1970). Habituation: A dual-process theory. Psycho-logical Review, 77(5):419–450.

Hearst, E. (1988). Fundamentals of learning and conditioning. In Atkinson, R. C., Herrnstein, R. J., Lindzey, G., and Luce, R. D., editors, Stevens’ Handbook of Experimental Psychology: Vol. 2. Learning and Cognition, pages 3–109. Wiley, New York, NY, 2 edition.

Hurvich, C. M. and Tsai, C.-L. (1989). Regression and time series model selection in small samples. Biometrika, 76(2):297–307.

Jorquera, O. E., Farfán, O. M., Galarce, S. N., Cancino, N. A., Matamala, P. D., and Vogel, E. H. (2025). A formal analysis of the standard operating processes (SOP) and multiple time scales (MTS) theories of habituation. Psychological Review, 132(6):1493.

Lewandowsky, S. (1995). Memory for the temporal spacing of events: Evidence for a single-process model. *Journal of Experimental Psychology: Learning*, Memory, and Cognition, 21(3):567–580.

McDiarmid, T. A., Bernardos, A. C., and Rankin, C. H. (2017). Habituation is altered in neuropsychiatric disorders—a comprehensive review with recommendations for experimental design and analysis. Neuroscience & Biobehavioral Reviews, 80:286–305.

Peeke, H. V. S. and Veno, A. (1973). Stimulus specificity of habituated aggression in the stickleback (Gasterosteus aculeatus). Behavioral Biology, 8(3):427–432.

Petrinovich, L. and Peeke, H. V. S. (1973). Habituation to territorial song in the white-crowned sparrow (Zonotrichia leucophrys). Behavioral Biology, 8(6):743–748.

Ponce, F. P., Vogel, E. H., and Wagner, A. R. (2015). The incremental stimulus intensity effect in the habituation of the eyeblink response in humans. Learning and Motivation, 52:60–68.

Rankin, C. H., Abrams, T., Barry, R. J., Bhatnagar, S., Clayton, D., Colombo, J., Coppola, G., Geyer, M. A., Glanzman, D. L., Marsland, S., McSweeney, F., Wilson, D. A., Wu, C.-F., and Thompson, R. F. (2009). Habituation revisited: An updated and revised description of the behavioral characteristics of habituation. Neurobiology of Learning and Memory, 92(2):135–138.

Rescorla, R. A. (1988). Behavioral studies of Pavlovian conditioning. Annual Review of Neuroscience, 11:329–352.

Rescorla, R. A. and Holland, P. C. (1976). Some behavioral approaches to the study of learning. In Rosenzweig, M. R. and Bennett, E. L., editors, Neural Mechanisms of Learning and Memory, pages 165–192. MIT Press, Cambridge, MA.

Thompson, R. F. (2009). Habituation: A history. Neurobiology of Learning and Memory, 92(2):127–134.

Thompson, R. F. and Spencer, W. A. (1966). Habituation: A model phenomenon for the study of neuronal substrates of behavior. Psychological Review, 73(1):16–43.

Wagner, A. R. and Vogel, E. H. (2010). Associative modulation of US processing: Implications for understanding of habituation. In Schmajuk, N. A., editor, Computational Models of Conditioning, pages 150–185. Cambridge University Press.

