## Supplementary material for "What group averages conceal: functional heterogeneity in human eyeblink habituation": Translational Abstract

### Public Significance Statement

Researchers commonly characterize the dynamics of habituation—the reduction in responding that occurs with repeated stimulus exposure—by fitting a curve to the group average and treating its shape as representative of the typical participant. This paper shows that the model selected for a group average can systematically differ from the model that best describes most individuals, even when the dataset is large and well-controlled. The difference arises because mathematically averaging curves of the same type can change the apparent shape of the resulting curve. This is therefore an effect of how aggregation works, not of the behavior itself. We provide a straightforward remedy: fitting candidate models to each participant individually and reporting the distribution of winning models alongside the group-average fit. This combined reporting practice can be implemented with standard statistical software and requires no additional data collection.
